# Dengue serotype-1 virus like particles induce antibody responses following HeLa cell expression

**DOI:** 10.64898/2026.04.08.717190

**Authors:** Smita Shrestha, Archana Maharjan, Roji Raut, Binod Manandhar, Binod Khadka, Ajit Poudel, Tek Raj Joshi, Dilip Chaurasia, Saugat R.C., Jarina Joshi, Rajani Malla, Lata Karki, Ram Prasad Aganja, Rajindra Napit, Krishna Das Manandhar

## Abstract

Dengue disease remains a significant global health threat, with current vaccines exhibiting variable efficacy and safety concerns. Virus-like particles (VLPs) offer a promising alternative by mimicking native virus structures without infectious genomes. We engineered a mammalian expression plasmid encoding Dengue-1 prM and E proteins, optimized for secretion using Japanese Encephalitis virus signal sequences, and transiently expressed it in HeLa cells. Purified VLPs exhibited spherical morphology (∼39 nm diameter) consistent with native virions, as confirmed by transmission electron microscopy. Immunization of mice with these VLPs elicited robust Dengue-1 specific IgG antibody responses. Our study demonstrates production of immunogenic Dengue-1 VLPs in HeLa cells, highlighting their potential as a vaccine candidate and a tool for serodiagnosis. Further characterization of VLP epitopes and protective efficacy is warranted to advance vaccine development.

**Importance:** Dengue remains a significant global health challenge, with serotype 1 being one of the dominant strains causing recurrent outbreaks in Nepal. Existing vaccines demonstrate limited efficacy and pose significant safety concerns, particularly in seronegative populations. To address these limitations, this study explores virus-like particles (VLPs) as a safer alternative vaccine platform. VLPs elicit robust immunogenicity by mimicking the structure of native virus while completely lacking genetic components. This study combines DENV1 structural proteins with optimized expression systems to enhance immunogenicity. This work is particularly significant as the first dengue vaccine research conducted in Nepal, directly addressing antigenic mismatches between existing commercial vaccines and locally circulating viral strains. Furthermore, the study provides scalable platform for developing region-specific dengue vaccines for other serotypes and flaviviruses.

## Background

Dengue is the fastest spreading vector-borne viral disease affecting humans [1]. It is an acute febrile illness with a wide range of clinical symptoms and severity, that ranges from asymptomatic cases, mild Dengue fever (also called classical Dengue fever) to life threatening severe infection Dengue hemorrhagic fever (DHF) and Dengue shock syndrome (DSS). About 28.5% of DHF progresses to Dengue shock syndrome (DSS) [2]. The mortality ranges between 1-20% depending on resources [3]. The viral hemorrhagic fever is considered the most serious biological illness, caused by pathogens such as Ebola virus, yellow-fever virus, lass virus, hantavirus and others [4]. Majority of hemorrhagic fever reported worldwide are related to dengue hemorrhagic fever making dengue virus one of the most serious threats to public health and to the global economy.

The dengue virus is transmitted to humans by the bite of infected female *Aedes* mosquitoes, predominantly *Aedes aegypti* and *Aedes albopictus*. In recent decades, the incidence of dengue has increased 30-fold due to the expansion of *Aedes spp*. habitat supported by climate change, globalization, urbanization, and resistance to commonly used insecticides such as DDT, and pyrethroids [5], [6]. New vector control strategies like deliberate introduction of *Wolbachia* endosymbiotic bacterium resulted in 95% reduction of dengue incidence however, its efficacy drastically reduced to 80% in parts of the world with high temperature, seasonal fluctuation, and high dengue endemicity [7], [8].

Vaccination is considered the safe and effective strategy to prevent from the dengue viruses that belong to *Flaviviridae* family of enveloped viruses; within genus *Flavivirus* carrying single stranded RNA genome. The virus consists of four antigenically distinct serotypes (DENV1 to DENV4) [9]. These serotypes share 25 to 40% sequence homology and 3 to 6% differences at amino acid level, giving rise to multiple genotypes [10], [11]. The genetic information of dengue is present on 10.7 kb long, single stranded, positive-sense RNA. The genome encodes a single polyprotein precursor. The single open reading frame of the genome is flanked by 5’ and 3’ untranslated regions (UTR) [12]. Inside the host cell, the precursor polyprotein is processed by viral and cellular proteases that generate three structural proteins: capsid (C), pre-membrane (prM) and envelope (E); and seven non-structural (NS) proteins: NS1, NS2A, NS2B, NS3, NS4A, NS4B and NS5 [13]. The central RNA genome is surrounded by a nucleocapsid core containing C protein. The virus is covered with dense icosahedral structure of envelope proteins which is lipid bilayer derived from endoplasmic reticulum of the host cell. Outside of envelope is embedded with E glycoproteins and M protein [12], [14]. The E glycoprotein exists as a homodimer, formed primarily with β-strands which folds into three structurally distinct components: Domain I, Domain II and Domain III, connected with flexible linkers. It mediates two crucial functions in the viral lifecycle: viral attachment and fusion with host cell [15].

The licensure of live attenuated tetravalent vaccines Dengvaxia®, and Qdenga® have given hope and opened new opportunities for the development of safe and efficacious vaccines against the above-mentioned structured dengue virus. However, it is important to acknowledge that these live attenuated vaccines carry risk of severe dengue in infants, increases replication or genetic reversion in immunocompromised individuals. Safety and efficacy data for both vaccines indicated varying effectiveness by age and dengue serostatus at the time of vaccination [16], [17]. There occurs to be higher degree of sequence similarity among members of Flaviviruses, resulting in similar genome organization, replication, and translation [18]. As a result, genes among flaviviruses can be interchanged to design a stable chimeric plasmid [18], [19]. This strategy was utilized by Dengvaxia to successfully produce chimeric live attenuated tetravalent dengue vaccine by replacing yellow fever virus (YFV) structural genes with dengue structural genes (prM and E) in the backbone of YFV [19], [20]. Hence, it is crucial to explore new avenues for the development of safe vaccines that provide balanced immunity against all serotype, and targets people of all age group, including dengue naïve individuals.

The virus-like particle (VLP) is a safer alternative vaccine approach as it triggers immune response without the presence of viral genome by mimicking the morphological structure of native infectious virus [21], [22], [23]. The lack of viral genome makes it safe as they are non-infectious. Another notable advantage of using VLP based vaccine against dengue is that the multivalent form, reduces the risk of Antibody-dependent enhancement (ADE) mediated severe dengue by eliciting balanced antibody response against all serotype [24].

Several dengue VLP vaccine has been developed based on prM and E protein using genes of native dengue virus [25], [26]. However, its *in-vitro* production has been reported to be quite low [27], [28]. The prM and E gene carried by Dengvaxia is of Asian Genotype I of DENV-2 (PUO-218 strain) [29] and its clinical trial failure in Thailand was partially linked to antigenic mismatch [30]. In Nepal, Dengue-1, Genotype I with strains from China and India is currently circulating [31], which is not incorporated in Dengvaxia [18], [32]. This antigenic mismatch between vaccine and circulating strain in Nepal could potentially lead to suboptimal protection and vaccination associated adverse effect, highlighting the urgent need to develop regionally tailored Dengue-1 vaccine.

To address these limitations, this study aims to improve the production of Dengue-1 VLP by combining natural sequence of Dengue-1 prM and E gene with signal sequence and transmembrane domain of JEV to enhance protein secretion and virus assembly. The DENV1 VLP developed in this study was evaluated for its ability to induce Dengue-1-specific IgG antibody response in mice, establishing baseline capacity for dengue vaccine research and development for the first time in Nepal where people are suffering every year.

## Results

### Transformed plasmid pDENV1 retained with promotor gene

The design of recombinant plasmid includes the origin of replication (*ori*), ampicillin resistance gene, CAG promoter region along with transcription enhancer, multiple cloning site containing insert of DENV1 complete *prM* and *E* gene, and Beta-globulin poly (A) transcription termination signal. The transcription of 2500 bp - DENV1 insert (*prM* and *E* gene) in pCAGGS plasmid was controlled by CAG early gene promoter/enhancer. The mRNA thus formed was terminated by beta-globin transcription terminator and stabilized by poly (A) signal [40]. The final length of the plasmid was 6,767 bp (Fig. 1).

**Figure 1.**
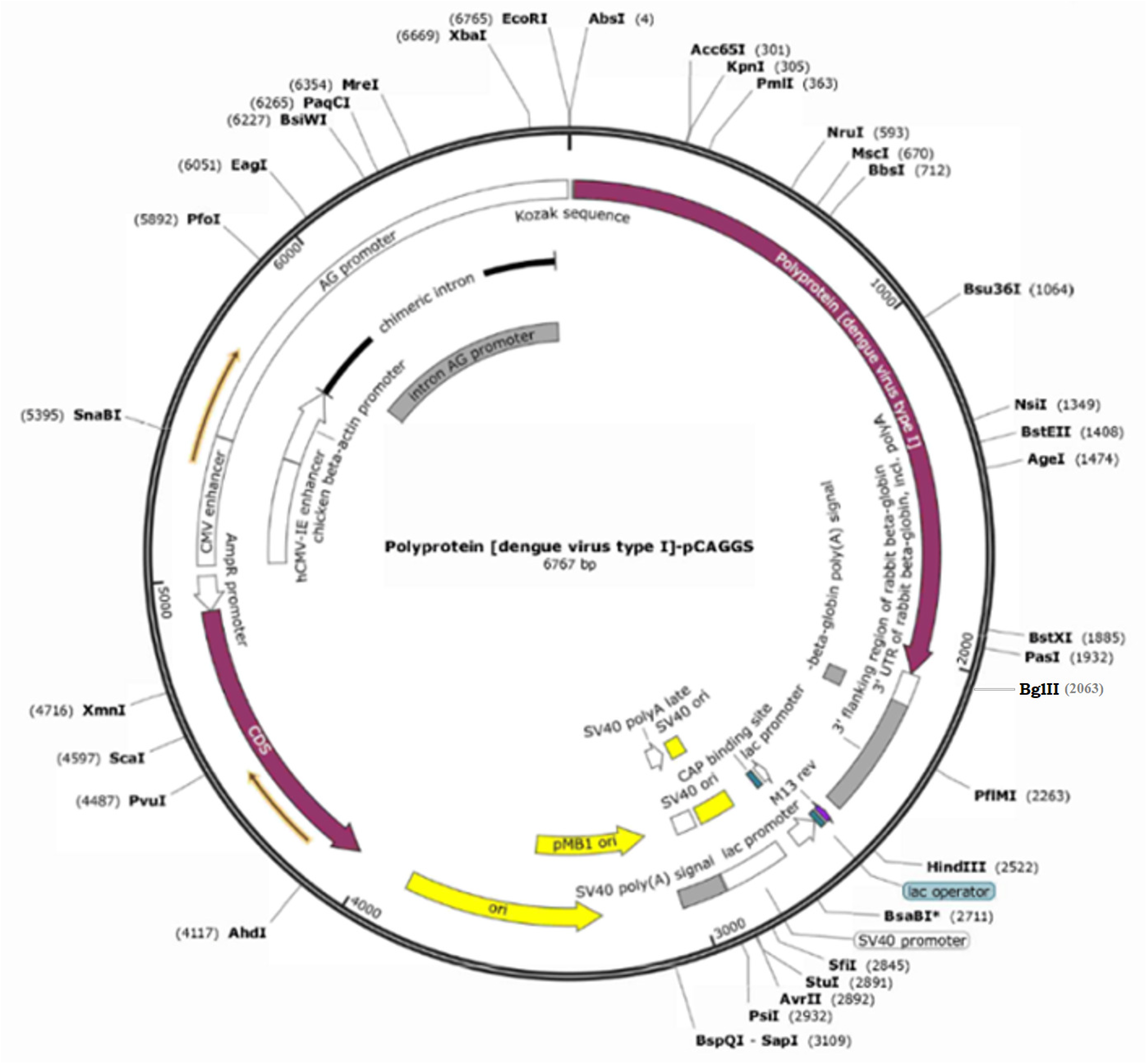
Schematic representation of DENV1 plasmid. The Dengue-1 prM andE gene is integrated into pCAGGS vector.

The successful addition of insert was confirmed by restriction enzyme digestion. The double digestion performed using EcoRI and BglII produced two fragments which were visible in gel electrophoresis image (Fig. 2.A). The linearized pCAGGS plasmid is represented by the larger 4000 bp fragment, while the smaller 2500 bp fragment represents the dengue insert drop out. These two bands, thus, validates, 2500 bp dengue insert was ligated in the plasmid.

**Figure 2.**
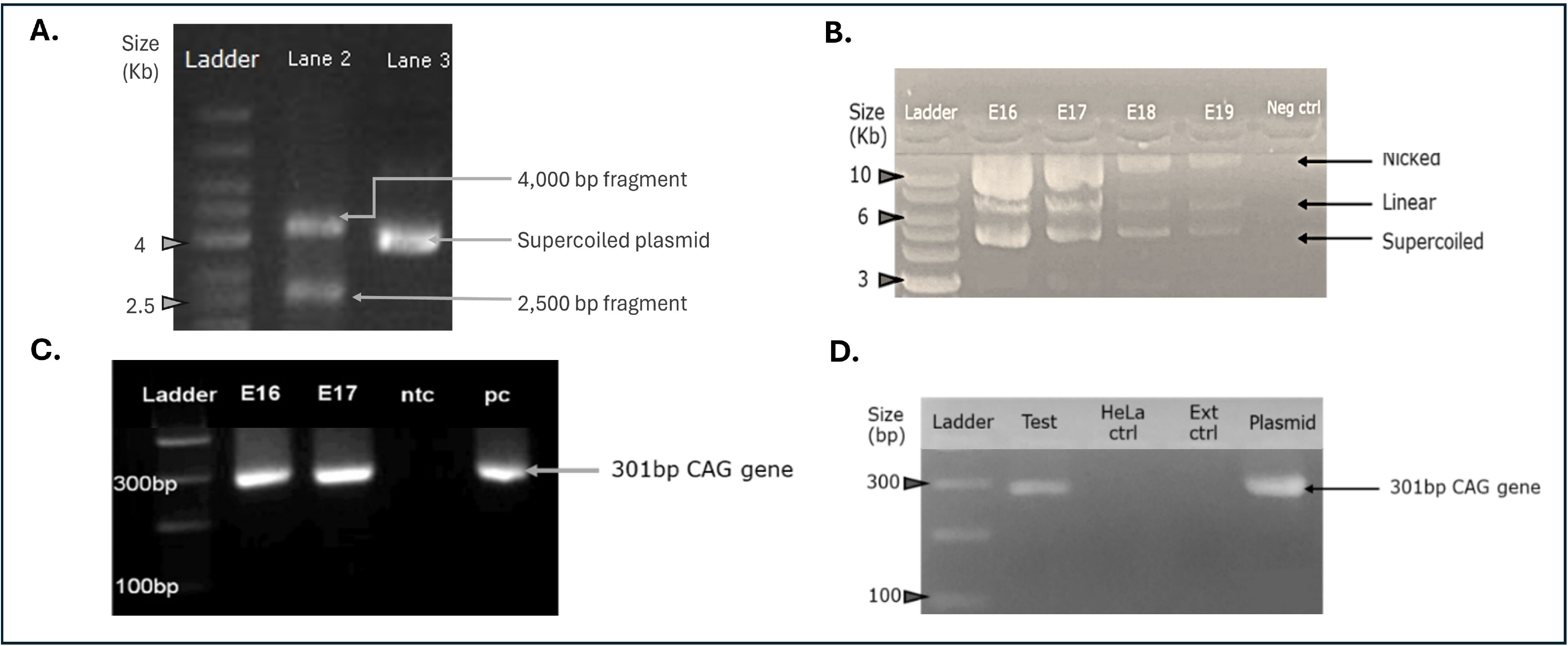
Gel electrophoresis images **A)** Plasmid DNA digested with enzymes. Lane 1-1kb DNA marker; Lane 2-plasmid after double digestion with ECoRI-BglII producing∼4,000 bp and ∼2,500 bp fragments; Lane 3-supercoiled plasmid. **B)** Agarose gel (0.8%) of plasmid DNA isolated from transformed DH5α. Lane1, 1kb DNA ladder; Lane 2 to 5, amplification of transformed colonies’ plasmid DNA with intense 3 plasmid forms: linear ∼7 kb, supercoiled ∼5 kb, nicked ∼10 kb; lane 6, negative transformation control. **C)** Agarose gel (1.5%) of CAG promoter gene amplification in transformed *E coli* culture by conventional PCR showing the intact of promotor gene in the propagated plasmid. Lane 1, DNA marker (100 bp); Lane 2-3, two transformed colonies’ plasmid amplifications; Lane 4, plasmid of transformation control (ntc); Lane 5, plasmid positive control (pc). **D)** Agarose (1.5%) gel of CAG promoter gene amplification in the transfected *HeLa* cell culture. Lane 1, DNA ladder (100 bp); Lane 2, transfected *HeLa* cell; Lane 3, transfection control; Lane 4, extraction control (Ext ctrl); Lane 5, plasmid used for transfection.

Propagation of DENV1 plasmid was achieved through transformation in *E. coli* DH5α. The transformed colonies containing DENV1 plasmid expressed intense bands at 5kb, ∼7kb and above 10kb corresponding to supercoiled, linear, and nicked open circular plasmids in gel electrophoresis respectively (Fig. 2.B). The size of the linear form of transformed plasmid corresponds with the designed *In-silico* form (Fig. 1).

To ascertain whether the extracted plasmid can drive protein expression and confirm the presence of plasmid, these plasmids were examined for the presence of CAG promoter. A positive amplification of 301 bp CAG gene was expressed in two (E16 and E17) transformed isolates (Fig 2.C).

### pDENV1 translated to DENV1 VLP in HeLa cell as confirmed by transient mRNA expression, SDS PAGE and TEM

The pDENV1 plasmids were employed for transient translation to DENV1-VLP in HeLa cells. Post transfected HeLa cells in 72 hours, were evaluated for changes in cell viability and morphology. The visible confluency and disintegration of live HeLa cells following transfection (test) dropped to approximately 50% (Fig. 3.B), compared to control non-transfected HeLa cells (Fig. 3. A). Trypan blue staining was performed to analyze the apparent morphological changes in both negative control culture and test cells (Fig. 3.C and D). Notably, the test cells exhibited visible protein assembly characterized by dense stains near the cell membrane, unlike the control cells where no such formations were observed. Transfection had visible impacts on the viability and morphology of HeLa cells.

**Figure 3.**
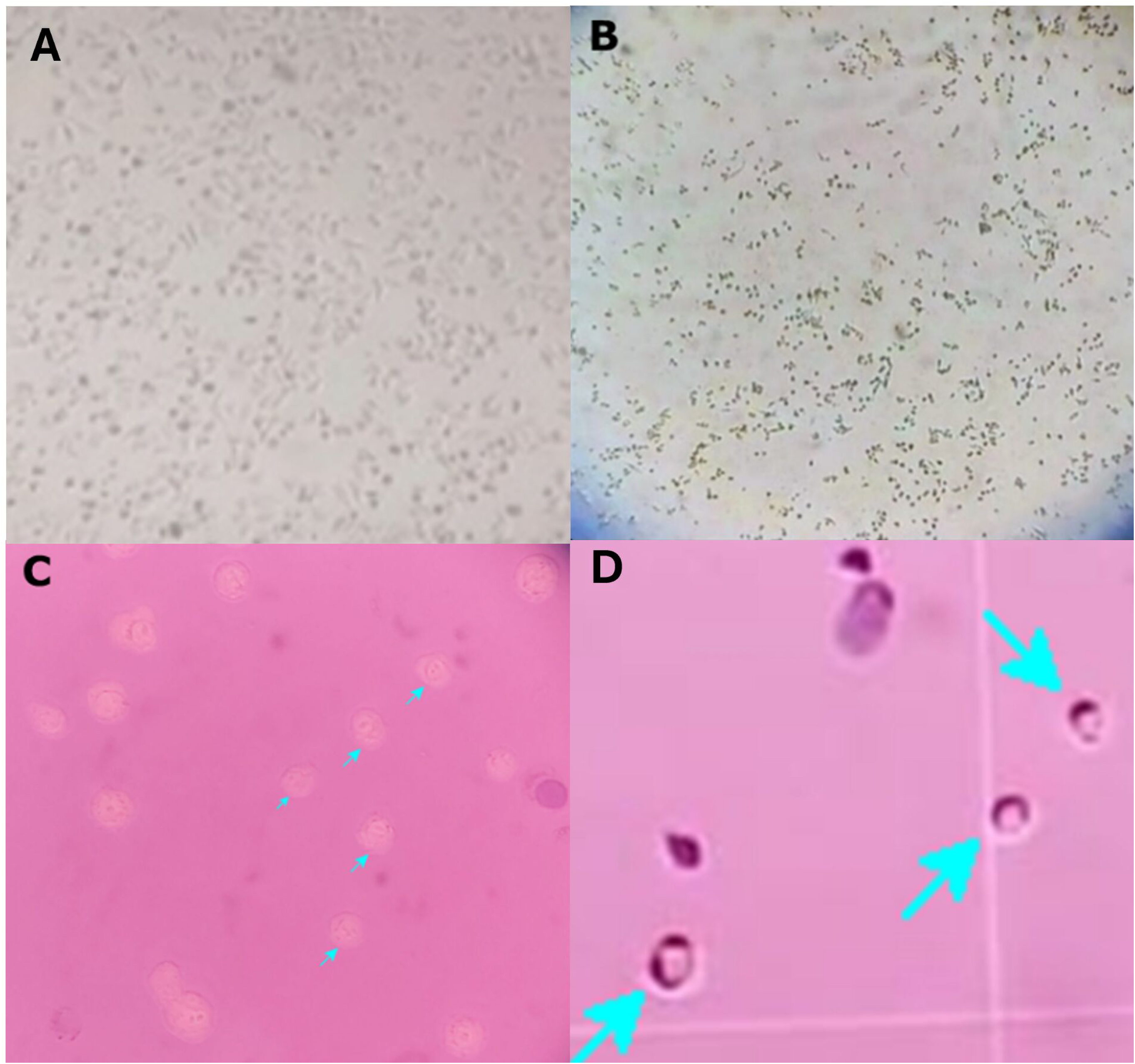
Microscopic analysis A) Control cell culture-Confluency 80% in 72 hrs. B) Post transfection- Confluency reduced to 50% in 72 hours. C) Morphology of non-transfected negative control subjected to trypan-blue stain at 40X microscopy. Arrowhead indicates live cells while stained are dead cells. D) Morphology of test cells observed at 40X microscopy subjected to trypan-blue stain. Arrowhead indicates the visible VLP protein accumulated on cell membrane.

Following RNA isolation from post transfected control and test HeLa cells, these cells were detected for the presence of transient mRNA containing DENV1 cassettes by semi quantitative PCR. Here, as well CAG promoter gene was amplified for the detection of DENV1 cassette. The 301 bp band consistently observed in transfected HeLa corresponds to the size of CAG gene in plasmid DNA (Fig. 2.D). Notably, the bands were visible only in RNA isolates of test cells. The same band was observed in plasmid utilized for transfection (Fig. 2C). No bands were observed in RNA isolates of control HeLa and extraction control. The test was duplicated, and the results were consistent.

A highly concentrated DENV protein band was observed in SDS PAGE gel at expected ∼60 KDa, consistent with the estimated molecular weight of dengue serotype 1 E-protein (Fig. 4). The total concentration of protein present in test supernatant fraction was higher (17.9 µg/mL) than of non-transformed negative control cells.

**Figure 4.**
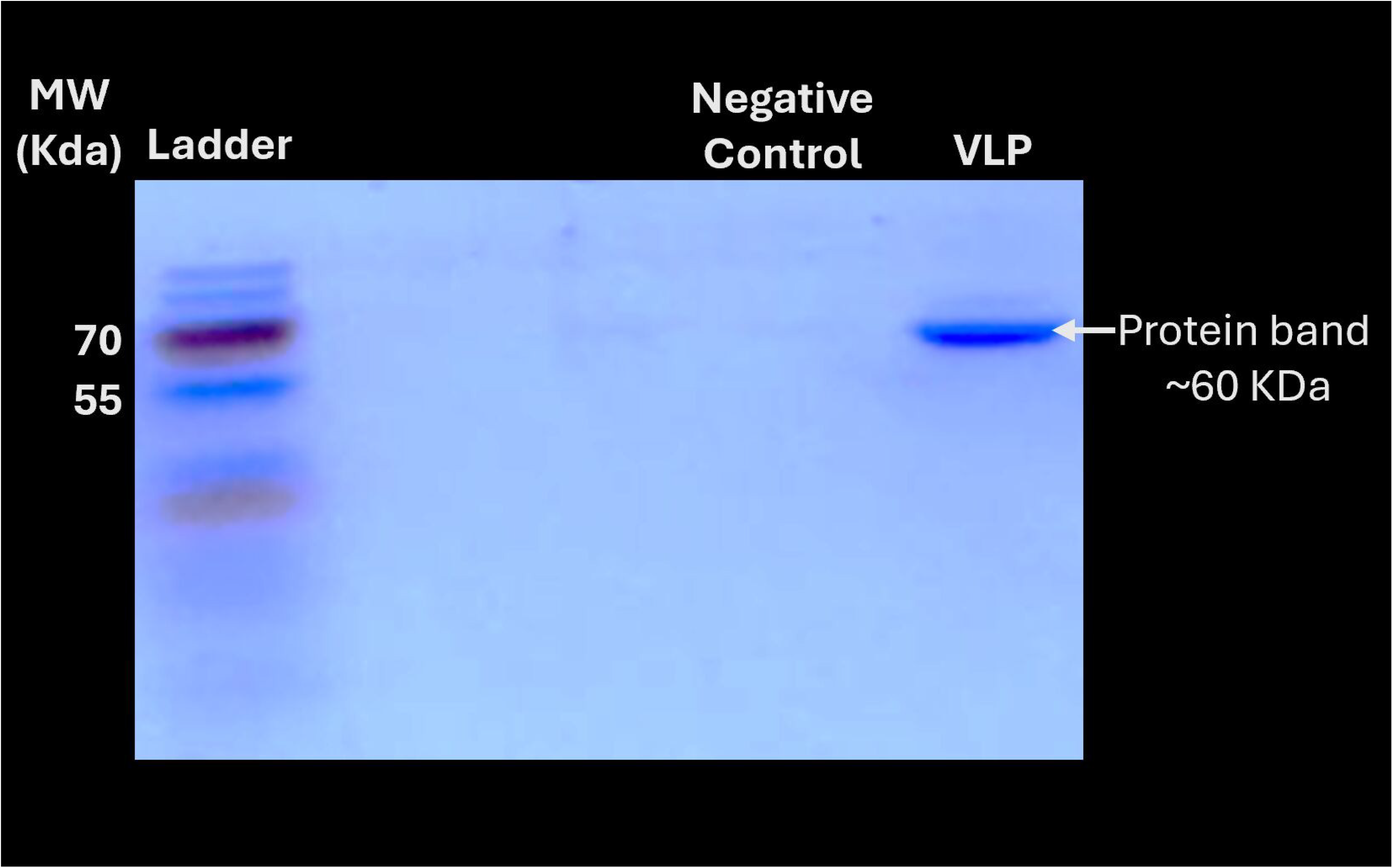
Estimation of molecular weight of purified VLP protein through SDS-PAGE. Lane 1- molecular weight marker; Lane 3- PEG purified culture supernatant of transfection control (Negative control); Lane 4- PEG purified culture supernatant of test. A highly concentrated DENV1 VLP protein band observed at 60 KDa.

The particle diameters (nm) obtained from TEM analysis showed variation between the control and VLP samples. In the control group, the mean particle diameter was 17.3 nm with a standard deviation of 7.2 nm and a 99% confidence interval of (15.22, 19.46) nm. In the VLP group, the mean particle diameter was 39.0 nm with a standard deviation of 18.6 nm and a 99% confidence interval of (31.07, 46.95) nm. The mean difference between VLP and control diameter size is 21.7 nm with a fold change of 2.2. Along with the VLPs, the fragmented VLP proteins were also observed in aggregated forms in the test samples which are not observed in the TEM picture of negative samples (Fig. 5).

**Figure 5.**
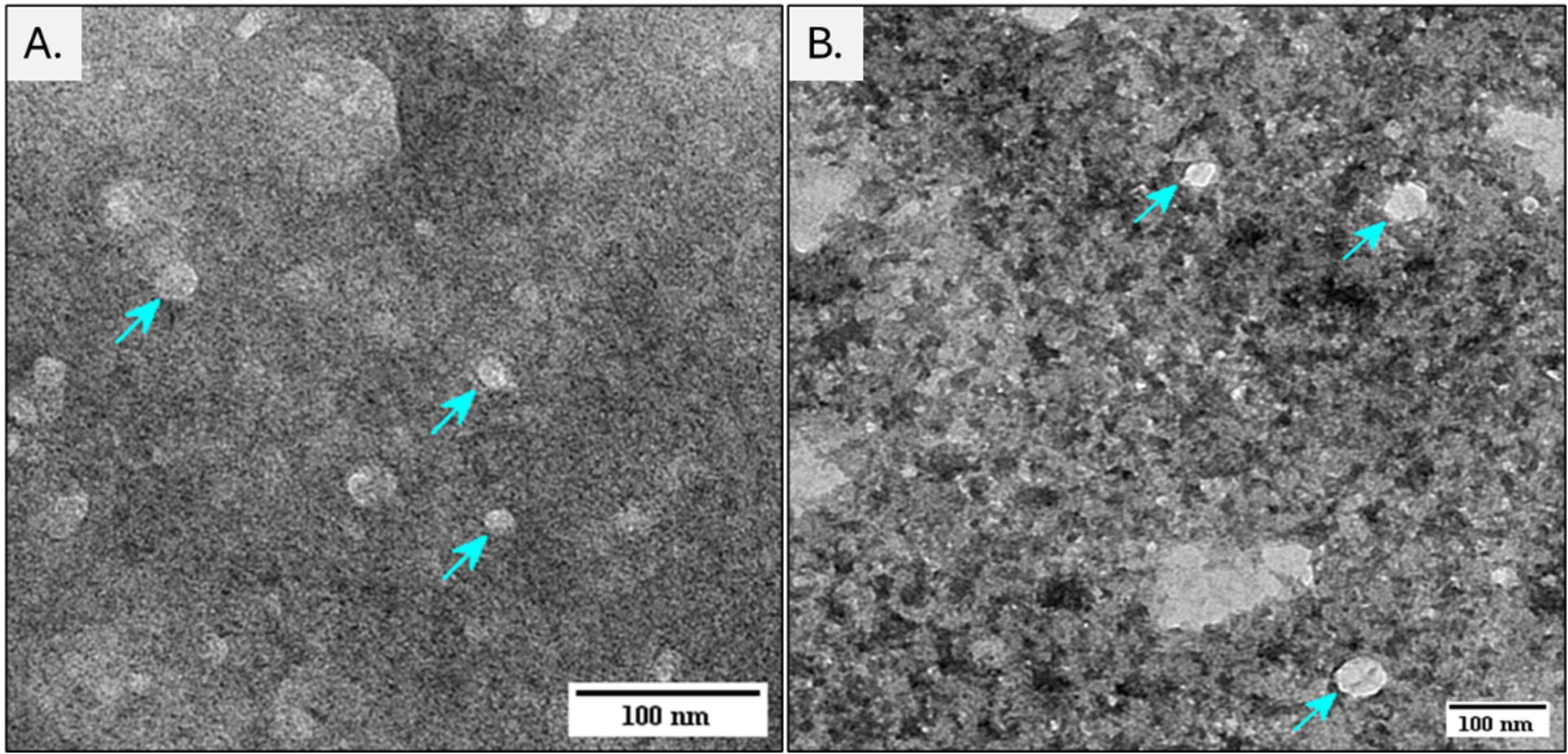
Electron micrographs of purified culture supernatant A) negative control (Scale bar 100 nm). Small particles observed are indicated by arrowhead. B) test supernatant for the characterization of particle shape and size (Scale bar 100 nm). A few spherical dense particles of size approx. 33 nm resembling the size of DENV-1 observed in test supernatant indicated by cyan arrowhead.

An independent Welch’s *t*-test demonstrated that the mean particle diameter of the VLP group was significantly greater than that of the control group. The right-tailed test yielded a *t*-value of 7.1 with 45.3 degrees of freedom and *p-value* < 0.00001. The 99% confidence interval for the mean difference in diameter between the VLP and control groups was (13.49–29.85) nm, indicating a significant increase in particle diameter in the VLP group compared with the control group (Fig. 6). Immunogenicity of DV1 protein The purified DENV1 protein was evaluated for its efficacy to elicit specific antibody response in female BALB/c mice. The immunized mice with purified DENV1 protein (50 µg/ml) and Freund’s adjuvant along with the first and second booster dose expressed specific anti-DENV1 IgG antibody which was evaluated by inhouse ELISA using purified DENV1 protein as the coating antigen. The ELISA reactivity of DENV1 immunized serum was significantly higher than control serum (Fig. 7).

**Figure 6.**
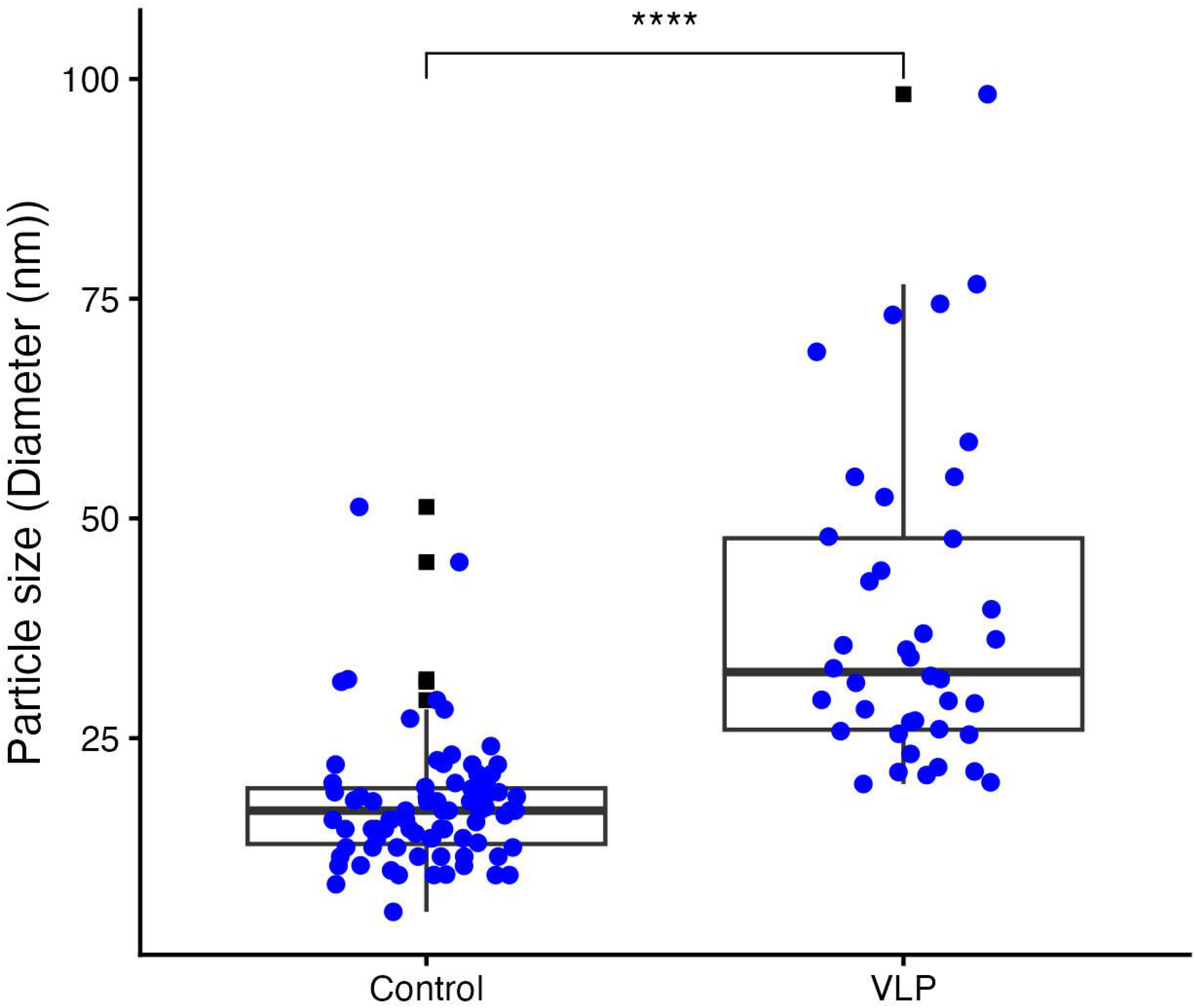
Comparative analysis of particle size distribution between negative control and test supernatant displayed by Box-and-whisker plot. Statistical significance was assessed by an independent right-tailed Welch t-test. An * indicates statistical significance (P<0.00001). *GraphPad Prism 8.0.2*

**Figure 7.**
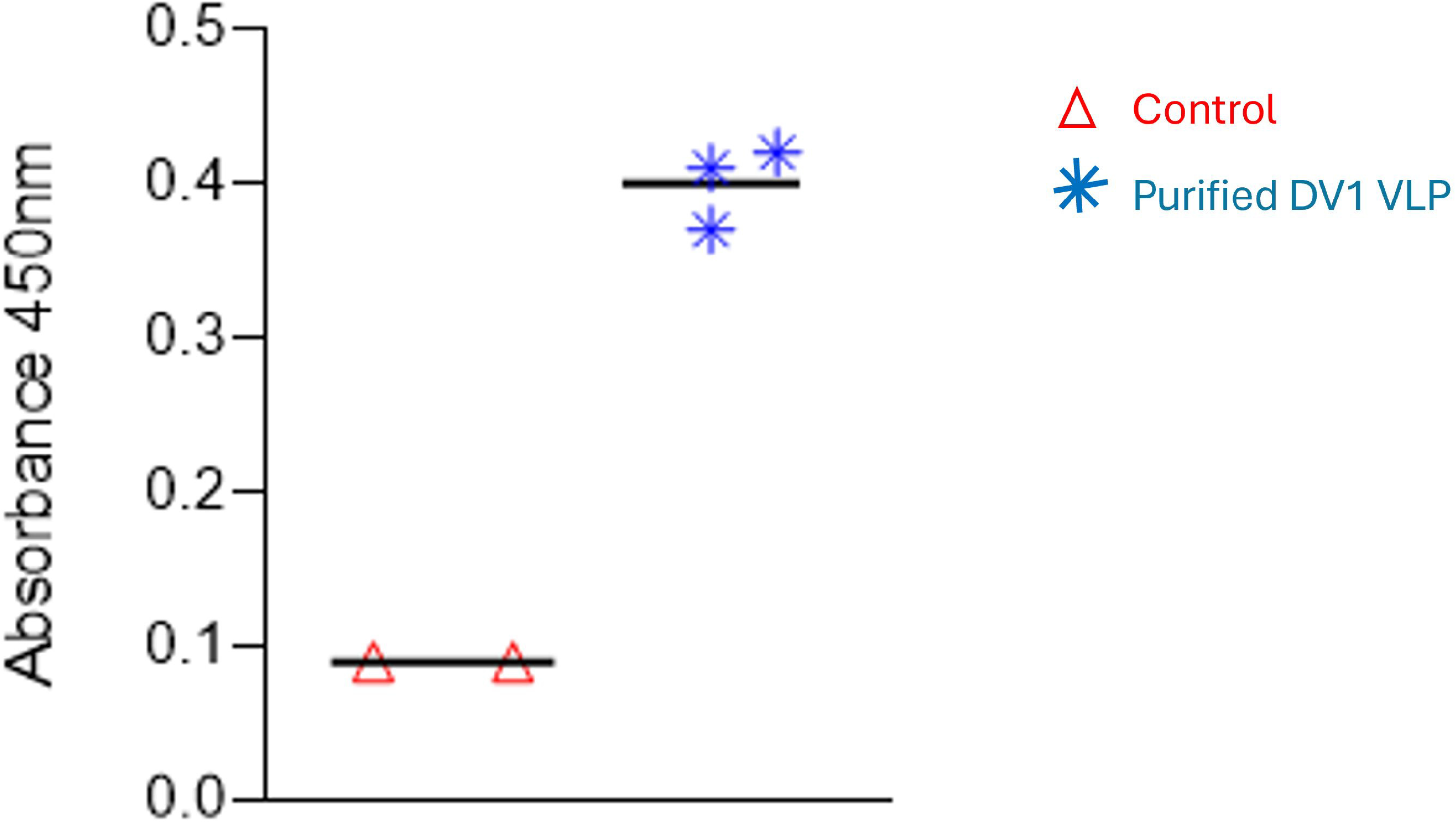
Immunogenic response of VLP compared to negative control as measured by ELISA. Serum antibody levels were quantified by measuring absorbance at 450 nm following in-house ELISA. The VLP immunized group showed significantly higher antibody response compared to the control group. Statistical significance was measured by an independent right-tailed Welch t-test (*p<0.01). *GraphPad Prism 8.0.2*

The control group exhibited low antibody levels, with a cutoff OD value 0.092 (3 times of mean control OD). In contrast, the VLP-immunized group demonstrated substantially higher absorbance, with a mean value of 0.40±0.026. The DENV1-immunized group showed at least 4.038-fold increase in anti-DENV1 IgG levels compared with the control group. An independent right-tailed Welch’s *t*-test showed significantly higher absorbance in the VLP group (*t-value* = 20.3, degrees of freedom = 2, *p-value* = 0.0012). The 99% confidence interval for the mean difference in absorbance was (0.158, 0.462).

## Discussion

Increasing prevalence of dengue and its gradual expansion into previously unaffected regions, even in face of climatic barriers underscore the pressing need for research on a secure and reliable dengue vaccine. The inefficiency of traditional live attenuated vaccines stresses the importance of investigating alternative vaccine approach. The VLP containing prM and E protein have been demonstrated by several groups as a potential dengue vaccine candidate [36], [41]. This study explores the feasibility of producing VLP that has a potential to be used as an alternate vaccine approach in the context of Nepal. In this study, a mammalian expression plasmid was engineered to produce DENV1 VLP in HeLa cells. These particles were extracellularly expressed. Upon purification, they were able to induce immunogenic response in mice after injection with second booster dose.

The Dengue VLP assembly and secretion is dependent on prM and E protein interaction. The strong ER retention signal located in transmembrane domain of E glycoprotein influences intracellular expression of VLPs [42], [43], which increases the steps for downstream processing. Several research groups have demonstrated the influences on the extracellular expression of Dengue-1 VLP with the use of signal sequence of JEV [44]. Clinical trial of the licensed JEV live attenuated vaccine which was constructed using the strain SA14-14-2 [18] has shown remarkable effectiveness, efficacy, and genetic stability in Asian countries including Nepal [45]. Therefore, signal sequence and transmembrane domain of JEV strain SA14-14-2 was selected in our study. Construction of the pCAGGS-DENV plasmid with JEV signal sequence along with Dengue-1 prM and E sequence was done to promote extracellular expression of VLP. The appearance of two distinct fragments after double digest with EcoRI/BglII validated the ligation of 2064 bp JEV-dengue fragment which was correctly oriented into the plasmid backbone (Fig. 2.A). Sequencing further confirmed the nucleotide level accuracy of the insert. This step was critical before proceeding to downstream applications, as it ensured that the expression cassette remained intact [46], [47].

Heterodimerized prM and E protein in endoplasmic reticulum is transferred to the Golgi complexes which pass on to the secretory vesicles through secretory pathways to be attached on plasma membrane before being released by budding. Staining pattern of DENV1 plasmid transfected cells revealed visible protein assembly near and around cell membrane (Fig. 3D) as compared to non-transfected cells (Fig. 3C) which validated the VLP trafficking destined to periphery of the cell for exocytosis. This indicates that the glycoprotein processing was correct as it marks the final stage of processing and translocation of proteins within the host cell, just prior to the budding of the VLPs. The protein assembly was formed during transcription and subsequent translation with virus specific information as encoded in gene cassette of our designed plasmid. This was confirmed by the detection of RNA transcripts in transformed cells (Fig. 2D). The overall yield of DENV1 VLP (17.9 µg/ml) confirmed the concentrated VLPs by PEG based purification.

TEM analysis revealed the VLPs were morphologically similar to natural DENV1 viruses. These VLPs appeared as monodispersed spherical particles, but their size was heterogenous (Fig. 5B). These purified VLP closely exhibited similar size and particle morphology to dengue VLP as reported in previous studies [36]. The particle size ranged from 20 to 98 nm, with average particle distribution at ∼39 nm. This variation in sizes could be attributed to the lack of dengue core structural proteins [48] and maturation difference between serotypes [41]. The observed particles on the negative control exhibited the typical features of extracellular vesicles which had well defined structure with smooth edges and uniform population size unlike the VLPs. Depending on the degree of cleavage and release of prM portion to form mature M protein, heterogenous population of immature, partially immature and mature dengue particles are released by host cell. As VLPs undergoes same maturation pathway as wild-type DENV; VLP produced by infected cells generate similar heterogenous population [49]. This could have possibly generated heterogenous VLP population of different diameters (Fig. 6). However, the extent of VLP maturation cannot be concluded in the study since the epitopes displayed by the VLP were not examined directly through western blot. Besides the size, the observed particles on the negative control exhibited the typical features of extracellular vesicles which had well defined structure with smooth edges and uniform population size unlike the VLPs. They are generated either through exocytosis or budding from the plasma membrane [50] and are involved in intracellular communication for cellular exchange of proteins, lipids and genetic material [51]

In our study, the presence of VLP in test supernatants was analyzed by comparing it with the supernatant of HeLa cells (used as negative control), unlike other studies that compared the size of engineered VLP with specific mutated virions. A direct comparison of the morphology of the VLP with the natural DENV1 virus was not conducted but the diameter of the VLP of our study was found to be comparable to mature dengue-1 virion as reported in other studies [48], [52].

The immunogenicity of the purified DENV1 VLP was well evaluated in BALB/c mice by measuring DENV1 specific IgG antibody. The absorbance value obtained from sera of DENV1 VLP immunized mice and from the control group indicated that all DENV1 VLP immunized mice had the absorbance reading above the threshold, suggesting a significant antigen-specific antibody response. The marked difference between the immunized and control groups suggests that the DENV1 VLP protein retains its antigenicity after purification and can elicit an antigen-specific antibody response. Freund’s adjuvant likely enhanced immunogenicity, as it is known to promote robust antibody production by sustaining antigen release and promoting Th2-type immune responses. The elevated IgG levels in the immunized mice demonstrate successful activation of B cells and subsequent antibody secretion in response to DENV1 VLP exposure.

These findings align with previous reports indicating that dengue virus proteins can serve as effective immunogens in murine models [8], [53], [54]. Importantly, the antibody response observed here suggests that the purified DENV1 protein is immunogenic in mice having potential to be a candidate vaccine with further investigation of neutralizing antibody production and diagnostic applications. However, to advance this candidature further, additional studies are needed to assess the neutralizing activity of the induced antibodies and to evaluate the protein’s protective efficacy in challenge models.

The study outlines a viable technique to produce DENV-1 VLP on mammalian cells utilizing structural proteins. The effective processing of proteins within the expression system was evident from the morphological characteristics of the particles, which closely mimicked the structural feature of mature dengue virus and elicited immunogenic response. Hence, a baseline for vaccine development capacity was initiated in Nepal with DENV1 VLP as potential vaccine candidate for the serotype. Additionally, it acts as a template for the development of vaccine against other serotypes and lays the groundwork for vaccine development within nation. It has potential to address the antigenic mismatch between vaccine and circulating strain in Nepal by incorporating E gene from dengue virus isolated in Nepal and could overcome the suboptimal protection and vaccination associated adverse effect.

## Methods

### Design of DENV1 Vector Plasmid

Dengue virus serotype 1 vector plasmid (pDENV1) was constructed with two major proteins: prM (Premembrane) and E (Envelope). The full-length sequence of DENV1 ectodomain prM and E protein was downloaded from NCBI database. The prM protein is encoded by *prM gene* (NCBI accession no. NP_733807.2) that encodes 166 amino acids. Similarly, the E protein, encoded by *E gene* (NCBI accession no. NP_059433.1) codes for 495 amino acids. These gene sequences were codon-optimized for better expression in HeLa cell line. The optimized nucleotide sequences of *prM* and *E gene* along with signal sequence and transmembrane domain of JEV strain SA14-14-2 were inserted in multiple cloning site (MCS) between *EcoRI* and *BglII* restriction sites of pCAGGS vector (GeneBank accession no. LT727518). The structure protein genes of Dengue-1 cloned into pCAGGS backbone was then custom-synthesized and made available by NovoPro synthesis. The *prM* and *E gene* were flanked by 5’ CAG promoter and 3’ transcription terminator respectively on pCAGGS vector. The size of pCAGGS plasmid was 4,267 bp and the size of the insert was 2,500 bp. The confirmation of the insert and quality of the plasmid was measured using next generation sequencing, and restriction digestion analysis.

### Transformation and plasmid extraction

The plasmid, pDENV1 was transformed by the heat-shock method into competent *Escherichia coli* DH5α. The competent DH5α cell was prepared following Hanahan procedure [33]. The transformed colonies were selected on nutrient agar (NA) plates, containing ampicillin (100 µg/mL). Single isolated colonies were selected and inoculated into 10 mL Luria broth (LB) containing ampicillin (100 µg/mL) for mass production and preservation, followed by validation with plasmid amplification and isolation. Plasmid was isolated using the alkaline-lysis method from an overnight culture of transformants grown on LB media containing ampicillin, which were incubated at 37° with constant agitation. This purified plasmid was subjected to agarose gel electrophoresis and visualized by UV transilluminator. The confirmation of transformed plasmid containing correct coding cassette of *prM* and *E gene* was detected using semi-quantitative PCR using forward primer CAGGS F (5’-TAATCAATTACGGGGTCATTAGTTCATAGC-3’) and reverse primer CAGGS R (5’-TCCCATAAGGTCATGTACTGGGCATAATGC-3’) [34].

### Propagation of DENV1 VLP

HeLa cell was transiently transfected with pDENV1 plasmid to evaluate the gene expression. Cell count of 0.3X10^6^ cells/well was seeded in 6 well culture plates to achieve 70% confluency. After 16 hours of seeding, these cells were transfected with 5µg of pDENV1 using Lipofectamine 2000 (cat. #11668030) at the ratio of 1:2 following the manufacturer’s instruction. The cell was then incubated at 37°C with 5% CO_2_ for 5 hours. The post-transfected cell was supplied with DMEM maintenance media containing 10% FBS and further incubated in similar condition for 72 hr to facilitate VLP production. The cells and culture supernatant were harvested separately 72 hr post transfection for VLP purification and characterization.

### Validation for transient transcription of DENV1 gene in cell culture

From the harvested cell, RNA extraction was performed by kit-based method. The cells suspended in PBS were lysed by boiling lysis method through brief incubation at 95°C. RNA was extracted using Qiagen RNA extraction kit (Qiagen, cat. #52904) as per the manufacturer’s instructions from the lysate.

The RNA was converted to cDNA using iScript cDNA synthesis kit (BIO-RAD cat. #1708891). Briefly, a 20 µL reaction mixture was prepared with 5µL RNA extracted template, 10 µL nuclease free water, 4 µL of 5X master mix and 1 µL Reverse Transcriptase enzyme. The reaction was primed at 25°C for 5 min, reverse transcribed at 46°C for 20 min, and terminated at 95°C for 1min. The resulting cDNA was utilized as a template for semi-quantitative amplification of the CAG promoter gene for validation of pDENV1 replication in mammalian cell line along with *prM* and *E* gene expression and translation to respective proteins.

### SDS-PAGE for validation of translated DENV1 VLP protein

The translated DENV1 prM and E proteins secreted in the culture were precipitated by polyethylene glycol (PEG) based precipitation using Polyethylene glycol (PEG-6000 and 2% NaCl) in equal volume. The mixture was incubated at 4°C on a constant shaking platform. After 48 hours, VLP was pelleted down using high-speed centrifugation platform, Optima XPN-100 ultracentrifuge (Beckman Coulter) at 10,000X g for 40 min at 4°C. The pellets obtained were then resuspended in 200µL of Sodium chloride-Magnesium sulfate buffer (SM buffer). The total protein concentration present in PEG-purified supernatant fractions was determined by Bradford assay (Thermo Scientific cat. #23200) according to the manufacturer’s instructions. A standard curve was generated using Bovine serum albumin (BSA), and protein concentration of unknown sample was determined by interpolation from the standard curve. Bradford reagent (1.0 mL) was mixed to all BSA standards and unknown samples followed by incubation at room temperature for 10 min to allow color development. Absorbance was measured at 595 nm using a microplate reader. All measurements were performed in duplicate, and the mean values were used for analysis.

The protein samples (150 ng) adjusted to 25 μL suspension with an equal volume of sample loading buffer and a 10 μL (Thermo Scientific cat. #26619) protein marker were loaded onto 5% stacking gel and separated on 12% resolving gel via electrophoresis set at 120 voltage for 90 minutes. After electrophoresis, the separated proteins were visualized by incubating the gel for 16 hrs. in Coomassie blue followed by overnight destaining of the gel. The separated protein bands in the SDS-PAGE stained with Coomassie blue were compared with the known marker band to confirm the protein by molecular weight.

### Transmission Electron Microscopy (TEM) for confirmation of translation and assembly of VLP

The morphology of translated assembled DENV1 VLPs were visualized by Transmission Electron Microscope (TEM). The purified VLP and control supernatant were fixed in 2% paraformaldehyde. Small droplets of fixed samples were placed in carbon coated copper grids. The grids were then stained with 2% saturated uranyl acetate and visualized at the TEM facility, AIIMS, Delhi. The micrographs were recorded by Talos (HR-TEM) operating at 200 kV. The diameter of the particles was measured by ImageJ software. A box-and-whisker plot was generated to visualize the particle sizes for the control and VLP groups. A right-tailed Welch t-test for independent and unequal variances was performed using GraphPad Prism 8.0.2 to compare the means.

### In vivo immunization and in vitro response to the VLP and the protein

Six female BALB/c mice (6-8 weeks old) were obtained from the Department of Plant Resources (DPR), Kathmandu, Nepal. Animals were housed under standard laboratory conditions. All procedures were conducted in accordance with institutional guidelines. Following acclimatization, mice were randomly divided into two groups (n=3 per group). The experimental group was immunized intramuscularly with DENV1 VLPs formulated with Freund’s complete adjuvant at equal ratio. While the control group received adjuvant alone. Each mouse in the experimental group received 50 μg of protein in the first and second weeks. Booster immunizations formulated with incomplete Freund’s adjuvant were administered twice at two-week intervals [35], [36], [37]. Blood samples were collected from the tail vein prior to the first immunization and at designated time points just before each inoculation. Sera were separated and stored at -80^0^C until further use.

The immunological responses expressed by mice against DENV1 VLPs were evaluated using sandwich ELISA. Assays were performed according to the manufacturer’s instructions using the InBios DENV Detect™ IgG ELISA kit [38] and with some modifications. Briefly, ELISA plate was coated with purified DENV1 VLP at 100 ng/well and incubated overnight at 4°C [39]. The plate was blocked with 100 µL blocking buffer for 1 hour at room temperature and washed with 1% PBS Tween-20. The test sera and standard controls were diluted 1:10 and 50 µL were added to the respective wells. Plates were incubated at 37°C for 1 hour followed by washing with 1X wash Buffer. Enzyme conjugated secondary antibody (50 µL/well) was added and incubated at 37 °C for 1 hour. After washing,150 µL of TMB substrate was added to each well and incubated at room temperature in the dark for 10 minutes. The enzymatic reaction was terminated by adding 50 µL of stop solution and incubating at room temperature for 1 minute. Absorbance was measured at 450 nm using a microplate ELISA reader (BMG Labtech, Germany).

### Statistical Analysis

A right-tailed independent Welch’s *t*-test was performed using GraphPad Prism 8.0.2 to determine whether the mean absorbance value of the VLP group was higher than that of the control group. *P* values less than 0.01 were considered statistically significant.

For the ELISA experiments, mice were assigned to both the control and VLP-immunized group. Antibody level absorbance values for each mouse were obtained.

For particle size analysis based on transmission electron microscopy (TEM), 80 particle diameter measurements were obtained for the control group and 40 measurements for the VLP group. Comparisons between the control and VLP groups were performed using an independent right-tailed Welch’s *t*-test to assess whether the mean values in the VLP group were significantly greater than those in the control group. Summary statistics, standard error, 99% confidence intervals for mean differences and mean fold changes were examined. Statistical significance was evaluated at the 1% significance level.

## List of abbreviations

ADE: Antibody dependent enhancement
C: Capsid protein
CAG: Chicken beta-actin promoter
DHF: Dengue hemorrhagic fever
DENV1 – 4: Dengue serotype 1, 2, 3, 4
DV1: Dengue Vector type-1 plasmid
DENV: Dengue virus
DNA: Deoxy-ribonucleic acid
E: Envelope protein
ELISA: Enzyme linked immunosorbent assay
JEV: Japanese Encephalitis
M: Membrane protein
NS: Non-structural proteins
PEG: Polyethylene glycol
PrM: Pre-membrane protein
RNA: Ribo nucleic acid
SDS-PAGE: Sodium dodecyl sulfate-polyacrylamide gel
TEM: Transmission electron microscopy
VLP: Virus like particle

## Declarations

### Author Contributions

Conceptualization-Rajindra Napit, Krishna Das Manandhar

Methodology-Rajindra Napit, Krishna Das Manandhar

Software-Rajindra Napit, Smita Shrestha

Validation-Krishna Das Manandhar, Rajindra Napit

Formal analysis-Krishna Das Manandhar, Rajindra Napit, Binod Manandhar

Investigation-Krishna Das Manandhar, Rajindra Napit, Smita Shrestha, Archana Maharjan

Resources-Krishna Das Manandhar

Data curation-Smita Shrestha, Archana Maharjan, Tek Raj Joshi, Dilip Chaurasia, Saugat RC, Binod Khadka, Lata Karki

Writing original draft preparation-Smita Shrestha, Archana Maharjan

Writing reviewing and editing-Krishna Das Manandhar, Rajindra Napit, Smita Shrestha, Ram Prasad Aganja

Visualization-Rajindra Napit, Smita Shrestha

Supervision-Krishna Das Manandhar, Rajindra Napit, Roji Raut

Project administration-Krishna Das Manandhar, Rajindra Napit, Ajit Poudel, Rajani Malla, Jarina Joshi

Funding acquisition-Krishna Das Manandhar, Rajindra Napit, Ajit Poudel, Rajani Malla, Jarina Joshi

### Ethics approval and consent to participate

The animal study protocol was approved by the International Review Board of Nepal Veterinary Council (protocol code Ethical/370 and date of approval May 05, 2025). Consent Statement is not applicable.

### Data Availability Statement

The raw data supporting the conclusions of the article will be made available by the authors on request.

## Acknowledgments

We are deeply indebted to Dr. Eans Tara Tuladhar, Dr. Aunji Pradhan, Ms. Lindsay Droit, Dr. William Telford, laboratory colleagues, faculty members, and staffs of the Central Department of Biotechnology (CDBT), Tribhuvan University for their unwavering support throughout this research.

## Consent for publication

The authors have reviewed and edited the output and take full responsibility for the content of this publication.

